# A novel approach to the functional classification of retinal ganglion cells

**DOI:** 10.1101/2021.05.09.443323

**Authors:** Gerrit Hilgen, Evgenia Kartsaki, Viktoriia Kartysh, Bruno Cessac, Evelyne Sernagor

## Abstract

Retinal neurons are remarkedly diverse based on structure, function and genetic identity. Classifying these cells is a challenging task, requiring multimodal methodology. Here, we introduce a novel approach for retinal ganglion cell (RGC) classification, based on pharmacogenetics combined with immunohistochemistry and large-scale retinal electrophysiology. Our novel strategy allows grouping of cells sharing gene expression and understanding how these cell classes respond to basic and complex visual scenes. Our approach consists of several consecutive steps. First, the spike firing frequency is increased in RGCs co-expressing a certain gene (*Scnn1a* or *Grik4*) using excitatory DREADDs (Designer Receptors Exclusively Activated by Designer Drugs) in order to single out activity originating specifically from these cells. Their spike location is then combined with *post hoc* immunostaining, to unequivocally characterize their anatomical and functional features. We grouped these isolated RGCs into multiple clusters based on spike train similarities. Using this novel approach. we were able to extend the pre-existing list of Grik4 expressing RGC types to a total of 8 and, for the first time, we provide a phenotypical description of 13 Scnn1a-expressing RGCs. The insights and methods gained here can guide not only RGC classification but neuronal classification challenges in other brain regions as well.

## Introduction

The retina contains two types of photoreceptors, rods for dim light and cones for daylight and colour vision. Furthermore, cone-contacting bipolar cells can be divided into ON and OFF types and further subdivided into more than a dozen different subpopulations (1). These parallel processed channels are further divided into a variety of functional output channels, so-called retinal ganglion cells (RGCs), which encode different features of the visual environment. There are ~1 million RGCs in humans and ~45,000 in mice (2,3), integrating the visual information processed from photoreceptors down the retinal neural network. Different types of RGCs extract very specific features from the visual scenery (4). This code is transmitted to postsynaptic targets in the brain, leading to visual perception. At present, more than 40 RGC types have been identified in the mouse retina (5,6). RGC classification is typically based on common anatomical features (7,8), responses to light (5,9–11) or on shared gene expression (6,12–14). Classification based on gene expression is relatively recent, and the majority of RGC groups sharing specific genes have not been phenotyped yet.

Current approaches for functional characterization of RGC subpopulations at pan-retinal scale are limited. Multielectrode arrays (MEAs) allow electrical recording from many RGCs simultaneously at high spatiotemporal resolution (15). Here we use a CMOS (complementary metal-oxide-semiconductor) MEA system consisting of 4,096 electrodes (2.67 x 2.67 mm arrays), allowing us to record light responses from hundreds to thousands of RGCs simultaneously at pan-retinal level and near cellular resolution (16,17). We selected two genes based on their sparse distribution across the RGC layer and their novelty for phenotypic characterization (Allen Mouse Brain Connectivity Atlas (2011)). *Grik4* (glutamate receptor, ionotropic, kainite subunit 4, HGNC: 4582) expressing RGCs have been partially described using a *Grik4* Cre mouse line (18). The other gene we investigated is *Scnn1a* (non-voltage gated sodium channel, epithelial 1 subunit alpha, HGNC:10599). Scnn1a Cre-induced recombination (*Scnn1a*-Tg3-Cre line) is present in sparse Layer 4 neurons, mostly in the somatosensory cortex (19). Current knowledge of Scnn1a expressing RGCs in the retina is limited to the fact that their dendritic arbour stratifies in sublaminas S1 and S2 (OFF layers) and in sublamina S4 (ON layer) of the inner plexiform layer (13). Here, we used the Cre-Lox recombination approach to specifically express DREADDs in these two Cre-lines to further investigate these RGC types.

Designer Receptors Exclusively Activated by Designer Drugs (DREADDs) (20) technology is a powerful new approach to pharmacologically dissect out the role of specific neuronal cell classes in network activity (21,22). DREADDs are an engineered version of muscarinic metabotropic receptors that allow precise control of G-protein signalling pathways. They are activated by “designer drugs” that have no endogenous receptors in the organism, such as clozapine-N-oxide (CNO). Most commonly used DREADDs are excitatory (hM3Dq, triggering release of calcium from organelles, leading to increase in intracellular concentration of free calcium and to membrane depolarization) In this study, we have generated Cre recombinase-mediated restricted expression of cell-specific DREADD (23) expression in either *Grik4* or *Scnn1a* reporter lines. We characterized the Grik4 and Scnn1a expression in RGCs and established an immunocytochemical atlas that is used to estimate the minimum cluster size for these cells. We have successfully isolated light-evoked responses in RGCs sharing either Scnn1a or Grik4 gene expression by combining excitatory DREADD activation, large-scale retinal CMOS MEA recordings and *post hoc* labelling of DREADD-expressing RGCs. We grouped the RGC responses into multiple clusters based on the similarity of the spike trains they generate in response to a series of stationary stimuli, thus extending or unravelling RGC types for the *Scnn1a* and *Grik4* gene pool.

## Results

To functionally validate RGC subgroups according to shared gene expression, we first established an immunocytochemical atlas of these cells. Building such a resource for Grik4 and Scnn1a expressing cells in the mouse retina is important to estimate RGC numbers and types in these two genetic pools. We used the intrinsic fluorescence signal of Grik4-DREADD (hereafter named Grik4) and Scnn1a-DREADD (hereafter named Scnn1a) cells to provide a detailed IHC map of Grik4 and Scnn1a expressing cells in retinal whole mounts and vertical sections. Each DREADD is tagged with hemagglutinin (HA) as well as mCitrine, allowing to visualize DREADD-expressing cells by immunofluorescence. We first investigated the distribution of Grik4 and Scnn1a cells in the ganglion cell layer (GCL) in retinal whole mounts using an antibody against Green Fluorescent Protein (GFP) to amplify the intrinsic mCitrine signal (Fig 1). Both lines exhibit sparse cellular distribution in the GCL, with Scnn1a cells significantly more abundant than Grik4 (Fig 1 A, B insets I and II). We calculated the cell densities in the GCL of three representative Grik4 (Fig 1 C) and Scnn1a (Fig 1 D) retinas. Grik4 and Scnn1a cell densities respectively vary between 10-25 and 40-100 cells/mm^2^. Both pools exhibit non-even distributions. Grik4 cells are more prominent in the dorsal-temporal periphery (Fig 1 C, yellow areas), while Scnn1a cells are more prominent in the ventral-nasal periphery (Fig 1 D, yellow areas).

**Figure 1:**
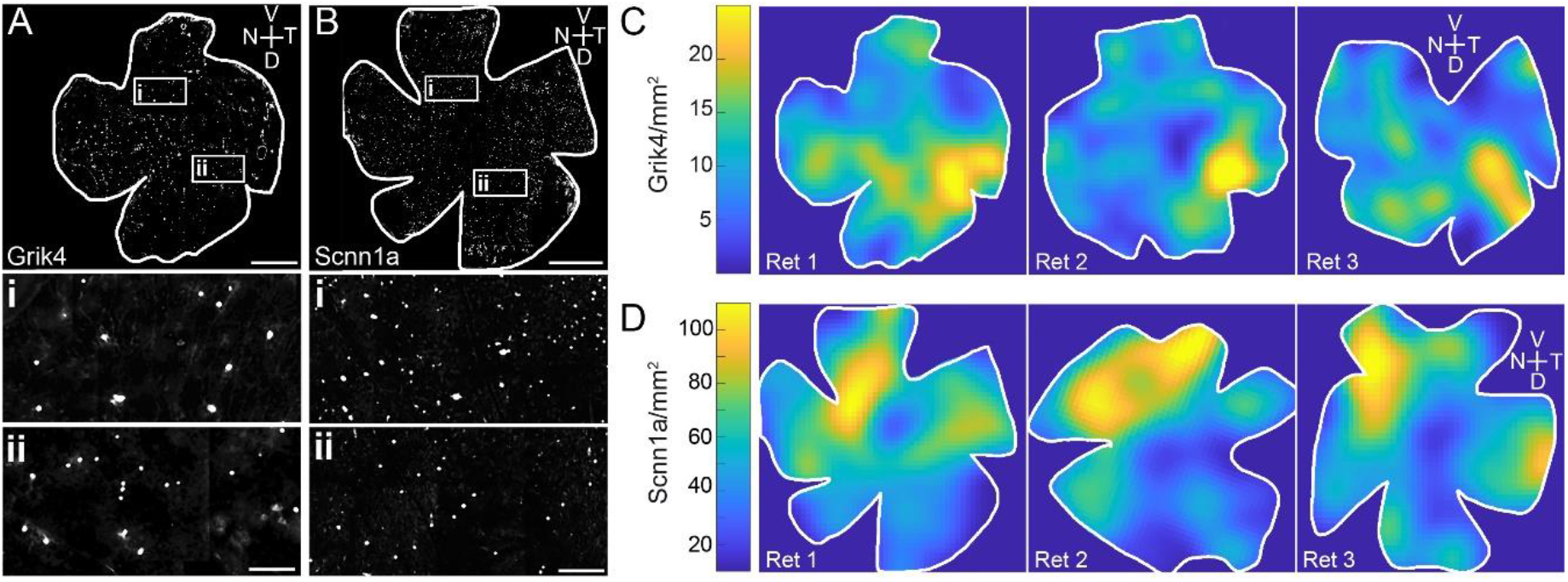
Grik4 and Scnn1a cells in the GCL are not homogeneously distributed. Whole mount antibody staining against Grik4 (A) and Scnn1a (B) DREADD GFP were imaged at the level of the GCL. All stained GFP cells were counted and the densities were calculated and presented in pseudocolors for 3 Grik4 (C) and Scnn1a (D) retinas. V= ventral, T = temporal, D = dorsal, N = nasal. Scale bar A, B = 1 mm; Scale bar A, B insets II = 100 μm.

To establish an immunohistochemical atlas of Scnn1a and Grik4 RGCs in the GCL, retinal whole mounts (Fig 2) and vertical sections (Fig 3) were stained for GFP, the calcium-binding protein marker parvalbumin (Fig. 2 A, B) (24), calretinin (Fig 3 A, B) (25), as well as a selective marker for RGCs in the mammalian retina (RBPMS (26) Fig 3 C, D)). In a first set of experiments (Fig 2 A, B) we labelled respectively against GFP (cyan) and parvalbumin (magenta) in retinal whole mounts of Grik4 (Fig 2 A) and Scnn1a (Fig 2 B) to find out whether there are cells in these two pools that co-express parvalbumin. The pool of parvalbumin RGCs in the mouse consists of 8-14 well-described RGCs sub-types (5,24,27–29). Quantification (n = 2 whole mount retinas) revealed that in Gri4 retinas, 6.6 ± 4.6 cells/mm^2^ cells are positive for parvalbumin in the central area, and 9.2 ± 11.2 cells/mm^2^ are positive in the periphery (i.e., approximately 50-65% of Grik4 cells are parvalbumin positive, Fig 2 A, bottom). For Scnn1a whole mounts (n = 2), we found 24.6 ± 10 parvalbumin positive cells/mm^2^ in the centre and 38.7 ± 13.6 cells/mm^2^ in the periphery (i.e., 40 - 50%). Given that the retinas were not additionally stained for RBPMS (both antibodies were raised in the same species), and some subclasses of displaced ACs are parvalbumin-immunoreactive, the real fraction of parvalbumin-positive RGCs expressing DREADDs could not be fully estimated. In summary, the number of DREADD GFP/parvalbumin cells is over 40% and higher in the periphery. NB the cardinal direction information was not noted for these experiments.

**Figure 2:**
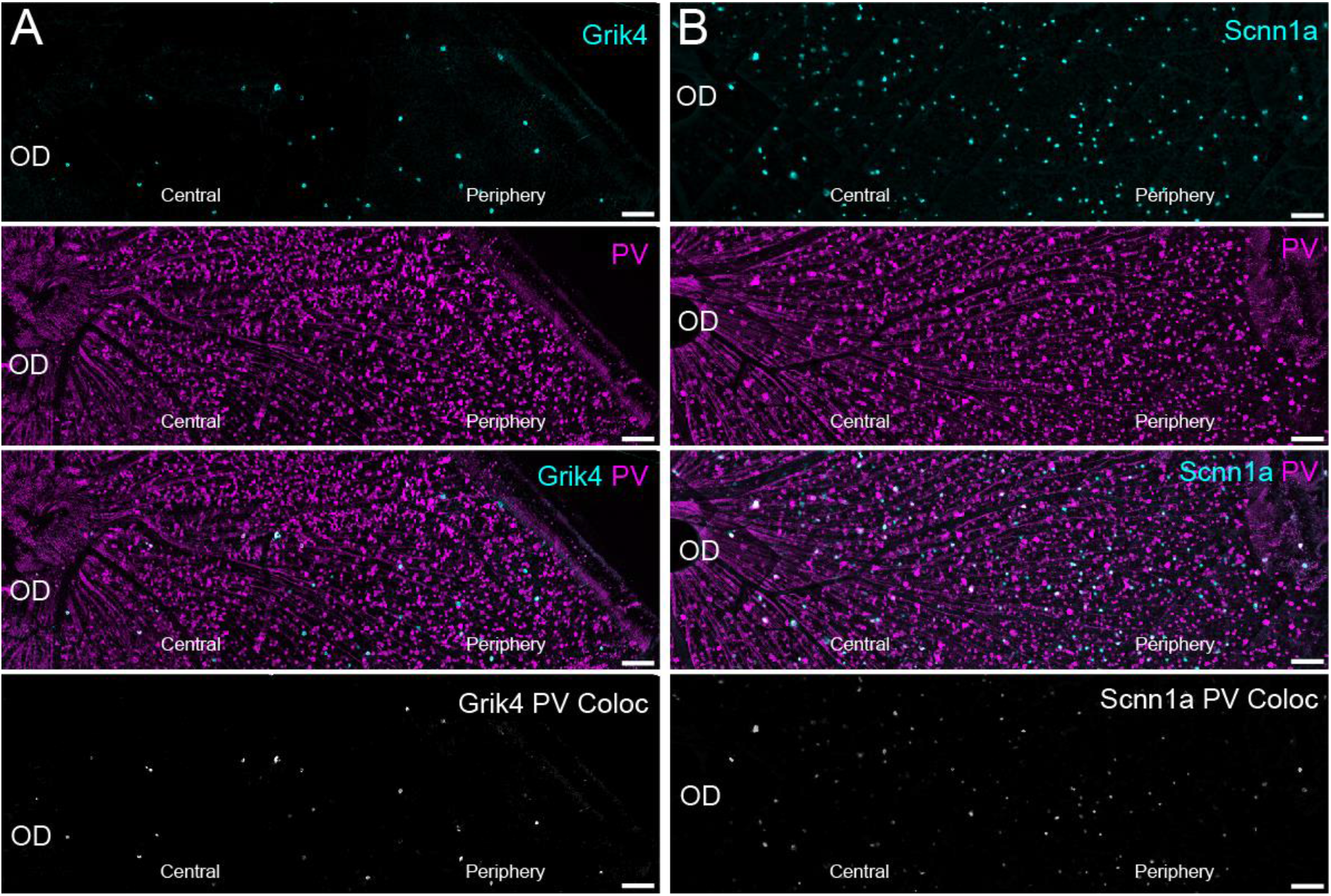
A large fraction of Grik4 and Scnn1a cells is Parvalbumin positive. Whole mount antibody staining against Grik4 (A) and Scnn1a (B) DREADD GFP in combination with Parvalbumin (PV, magenta) revealed that a large fraction of the DREADD cells in the ganglion cell layer is expressing also Parvalbumin (middle and bottom). Bottom images show DREADD and PV image layers multiplied, thus revealing colocalised cells. Scale bars = 100 μm.

**Figure 3:**
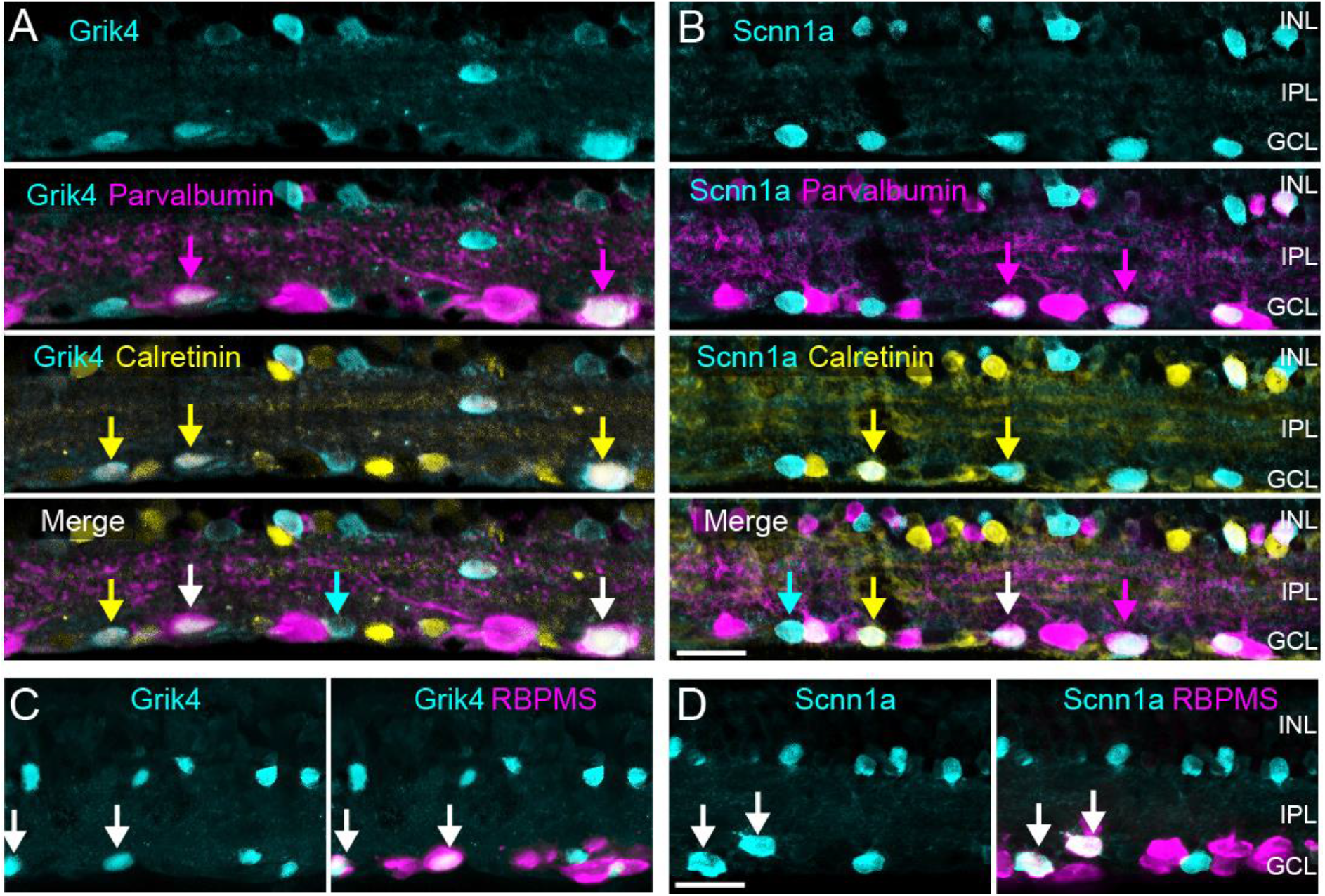
Grik4 and Scnn1a DREADD are expressed in multiple RGC types. Vertical sections of Grik4 and Scnn1a retinas were triple stained A, B) for GFP (cyan), Parvalbumin (magenta) and Calretinin (yellow) or C, D) GFP (cyan) and RBPMS (magenta). INL = inner nuclear layer, IPL = inner plexiform layer, GCL = ganglion cell layer. Scale bar in B, D = 20 μm.

Our initial findings indicate that a large fraction of DREADD cells in the periphery of the GCL are parvalbumin positive. The next experiments thus aimed to find out whether these DREADD-expressing cells can be further subdivided into smaller groups. Besides parvalbumin, the expression of calretinin, another calcium-binding protein, is well described in RGCs (25). Hence, we triple-labelled vertical sections (Fig 3 A, B) for GFP (cyan), calretinin (yellow) and parvalbumin (magenta). We found that the population of Grik4 (Fig 3 A) and Scnn1a (Fig 3 B) expressing cells (cyan) in the GCL consists of at least three different types of cells: GFP, GFP/calretinin and GFP/calretinin/parvalbumin (Fig 3 A, B arrows). Moreover, for Scnn1a we found an additional subset of GFP/parvalbumin cells (Fig 3 B magenta arrows). Thus, some of our Grik4 and Scnn1a RGCs should overlap with the known 8-14 parvalbumin RGC types and the ~10 different calretinin RGC types (25). So far nothing is known about Grik4 or Scnn1a expression in parvalbumin and calretinin RGCs. However, the rare parvalbumin/calretinin co-expressing RGCs have been functionally described before (27,30,31), hence their typical response characteristics should be easy to spot in our Grik4 and Scnn1a functional clusters (see below).

The GCL consists mainly of RGCs but it also contains displaced amacrine cells (dACs). We investigated whether dACs contribute to the pool of Grik4 and Scnn1a cells in the GCL. First, we double-labelled Grik4 and Scnn1a retinal vertical sections against GFP (Fig 3 C, D cyan) and RBPMS (Fig 3 C, D magenta). Approximately 50% of the GFP labelled Grik4 cells in the GCL are RBPMS positive (Fig 3 C, arrows), and the other cells presumably are dACs as they did not stain for RBPMS. In support, the somata of putative dACs in the GCL are relatively small, a key feature of these cells (Fig 3 C, GCL). A similar pattern was found for Scnn1a cells in the GCL but here, the majority (50-60%) of Scnn1a cells are RGCs (Fig 3 D, arrows). Sparsely distributed cells expressing either Grik4 (Fig 3 C, cyan) or Scnn1a (Fig 3 D, cyan) were present in the proximal inner nuclear layer (INL). These cells did not stain for RBPMS, confirming they are amacrine cells (ACs).

In summary, the pool of Grik4 and Scnn1a cells in the GCL respectively consists of at least three and four different cell types. It is likely that these different cell types reflect RGCs and not dACs because we did not find any putative Grik4- or Scnn1a-positive ACs in the INL expressing calretinin and/or parvalbumin (Fig 3 A, B, INL). Such information is important for the validation of our novel approach for the functional classification of these same RGCs. If our cluster size falls below these numbers, it would indicate that our method is flawed.

The presence of Grik4 and Scnn1a DREADD ACs was not expected. Excitatory Grik4 and Scnn1a DREADD ACs will have profound effects on their postsynaptic RGC partners. Indeed, a single AC affects nearly every RGC within reach of its dendrites (32). For example, let us consider a simple network with one population of ACs which modulate one population of RGCs (Fig 4 A). Figure 4 explores different scenarios. If DREADDs are expressed only on RGCs (Fig 4 A, scenario 1), adding CNO will have a direct excitatory effect on RGCs, increasing their activity and straightforward isolation for further classification. On the other hand, if there are DREADDs on ACs as well, CNO will either depolarise only DREADD ACs (Fig 4, scenario 2) or simultaneously DREADD ACs and RGCs (Fig 4A, scenario 3). These situations are more challenging, as they make it difficult to identify DREADD RGCs based solely on activity levels. Depending on whether they are excitatory or inhibitory cells, such DREADD ACs would lead to increased or decreased activity in non-DREADD RGCS (scenario 2), thus leading to false-positive identification of DREADD RGCs. Finally, the presence of DREADDs both in ACs and in RGCs will lead to an uncertain outcome, depending on the nature of the AC (Fig 4 A, scenario 3).

**Figure 4:**
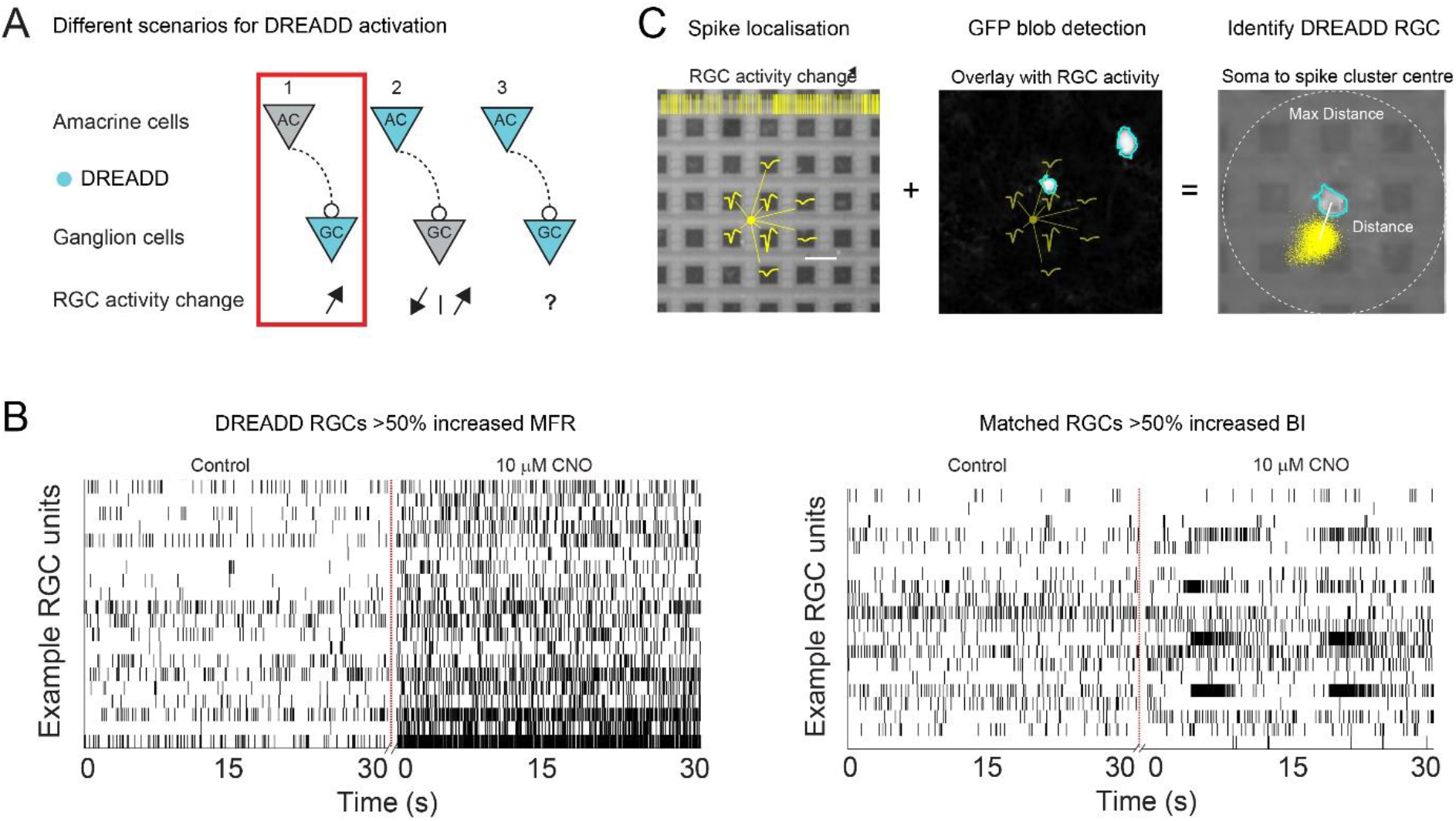
Registering GFP RGCs with nearby isolated spike centre clusters to unequivocally isolate DREADD RGCs. Activation of DREADD ACs can lead to different scenarios in RGC activities (A). DREADDs in RGCs can be activated with Clozapine N-oxide (CNO) and lead to an increase in firing frequency (B, left) and sometimes also in bursting activity (B, right). Vertical lines represent spike times plotted as a function of time. The interpolation of spike amplitudes from the same RGC defines the potential electrical source location of the isolated RGCs (C, left). Micrographs of DREADD-GFP expressing cells in the GCL were aligned with the MEA grid (C, middle). Spike centres from RGCs that showed an increase in activity and near GFP labelled RGCs were registered as potential DREADD-expressing RGCs (C, right).

Whatever the scenario is, we cannot rely on changes in DREADD RGC activity alone to identify DREADD-expressing RGCs. Hence, we applied a two-steps protocol to unequivocally isolate DREADD RGCs. In the first step, we pre-identify cells that show either an increase in spontaneous firing rate (Fig 4 B, left) or in the Burst Index (Fig 4 B, right) (33) in the presence of CNO. Next, the physical position of these identified RGCs is correlated with micrographs of DREADD-GFP expressing cells in the GCL (Fig 4 C). Further technical details and codes are available in the Method section and our GitHub documentation. Briefly, due to the high electrode density, the activity of one RGC is recorded on multiple adjacent electrodes (Fig 4 C, left). The x and y position of the spike origin (Fig 4 C, left, yellow dot) is calculated using the spike interpolation algorithm in Herdingspikes2 (34,35). A stitched high-resolution image of GFP stained cells in the GCL is then aligned with the MEA electrode grid (Methods & GitHub documentation). Image segmentation techniques are used to threshold the GFP foreground from the background and a blob detection is applied to detect the outlines and centre of Grik4 and Scnn1a GFP positive cells in the GCL (Fig 4 C, middle). Lastly, the Euclidian distance between the centre of the GFP cell and the spike origins of the surrounding RGCs is calculated. RGC units showing >50% increase in activity levels AND localised within a 60 μm radius of the GFP centre (Fig 4 C, right) are registered to be DREADD RGCs for Grik4 and Scnn1a retinas, respectively.

The rationale for choosing a 50% activity increase threshold and 60 μm distance is as follows. Spikes originate from the axon initial segment (AIS) rather than from the cell body. Therefore, we expect the spike cluster centre to be slightly eccentric with respect to the soma itself. The exact location of the AIS in RGCs can be very close to the soma (<30 μm), or sometimes rather distant along the axon (>30 μm), depending on the eccentricity of the cell with regard to the optic nerve head (36,37). For every detected Grik4 (Fig 5 A) and Scnn1a (Fig 5 B) GFP cell, we collected the activity change information (spiking rate or burst index change, depending on what is more prominent) from all RGCs after application of CNO within a 200 μm radius (Fig 5 A, B). Most cells do not express DREADDS, and therefore were not or only very slightly affected by CNO, showing a peak around 0% RGC activity change (Fig 5 A, B, magenta distribution curve). The RGC activity change curve (magenta) falls back to baseline levels at around 100% RGC activity change both for Grik4 (Fig 5 A) and Scnn1a (Fig 5 B). We defined the 50% full width at half maximum (FWHM, Fig 5 A, B dashed line) value of the curve (Fig 5 A, B) as a good threshold for defining spiking (and bursting) activity changes in DREADD RGCs after application of CNO (Fig 4 B). Notably, higher RGC activity changes (>50%) were closer (<30 μm) to the soma (Fig 5 A, red), further confirming that these activity changes belong to the corresponding DREADD GFP cells. A similar relation between “distance from soma” vs “RGC activity change” can be found in Scnn1a retinas (Fig 5 B, red) although some potential DREADD RGCs appear to be located further away from the soma (up to 100 μm) than Grik4 cells. Both red distribution curves (Fig 5 A, B) have their peak around 10 μm and plateau around 60 and 120 μm, respectively for potential Grik4 and Scnn1a RGCs. The FWHM distance from soma value for Grik4 is 30 μm and for Scnn1a, 60 μm. Based on these observations, we concluded that using the 60 μm radius was sufficient to reliably determine whether an isolated spike cluster corresponds to a specific GFP-expressing cell and thus represents a DREADD RGC. It is also close to the maximum thresholds given in the literature (36,37).

**Figure 5:**
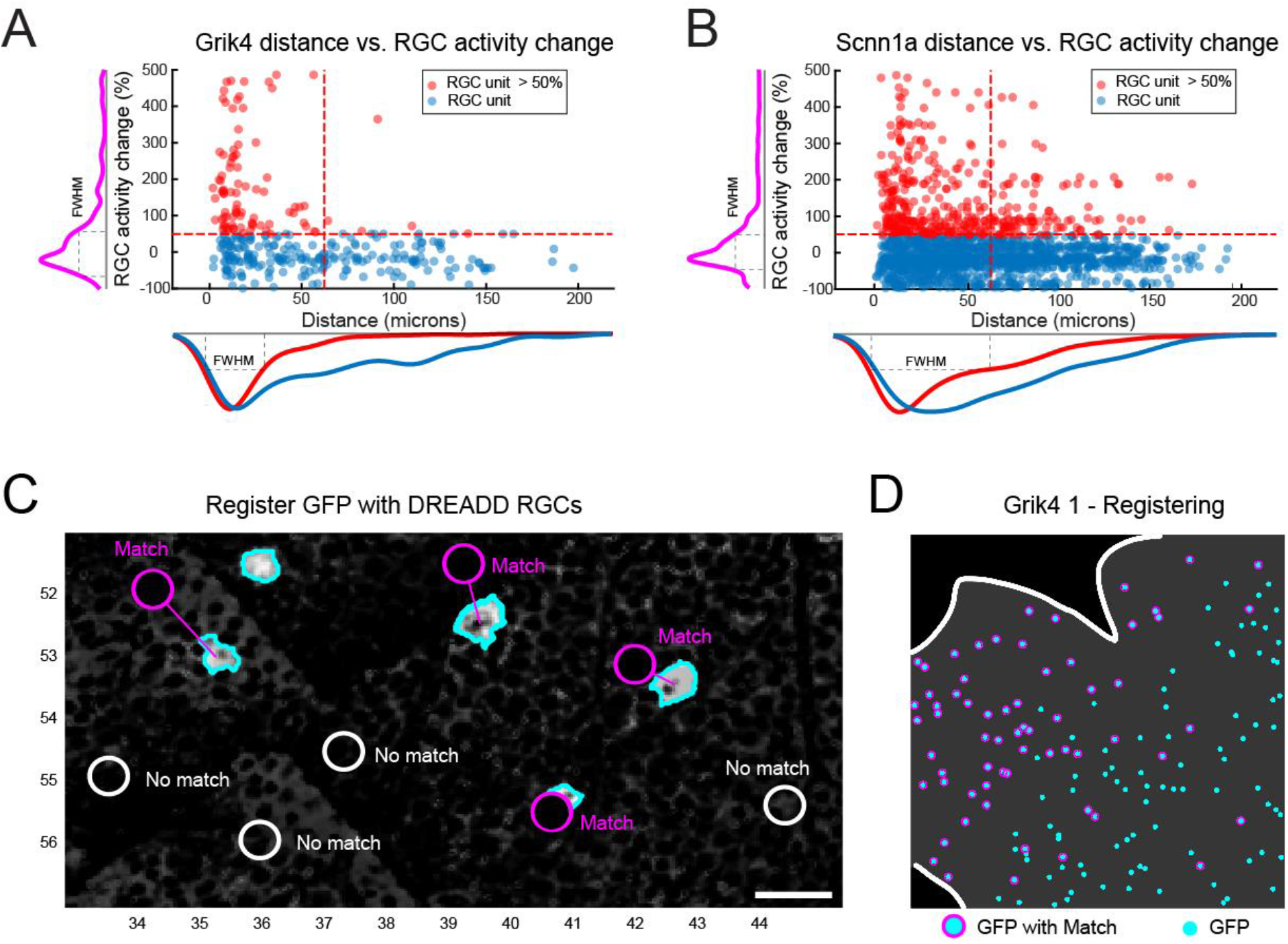
Defining suitable thresholds to isolate DREADD RGCs. The Euclidian distances from all spike cluster centres within a radius of 200 □m to a Grik4 (A) or Scnn1a (B) GFP cell are plotted as a function of their RGC spiking/bursting change (higher value). Red dots represent RGCs clusters that exceeded the 50% RGC activity change threshold. The y-axis distribution curve is summarising blue and red dots (magenta). The x-axis distribution curve is separating red and blue dots. FWHM = full width at half maximum. C) RGC units that were physically located within a 60 □m radius AND exhibited at least 50% increase in spiking or bursting rate were defined as DREADD RGCs (Match). White circles represent RGC units that did fall into the set criteria (No match). D) The steps are applied to all GFP cells. NB the recording area is generally not completely covering the entire 64 x 64 array i.e., covering all GFP cells.

The process of correlating physical spike positions with structural imaging allowed us to unequivocally isolate Grik4 and Scnn1a RGCs. Such cases are referred to as “Match”, and those falling outside these boundaries are classified as “No Match” (Fig 5 C). We successfully analysed three Grik4 and five Scnn1a retinas to register “Match” (Fig 5 C, D, magenta) cells. Figure 5 D is a representative example from one of the Grik4 retinas. Note that the recording area is generally not completely covering the entire 64 x 64 array. In total, we identified 82 “Match” RGCs in the three Grik4 retinas and 107 in the five Scnn1a retinas. However, most RGCs did exhibit changes (>50%) in firing rate or bursting without revealing any GFP signals, hence they are unlikely to be Grik4 or Scnn1a RGCs. These cells are most likely other RGC types affected by DREADD expressing ACs, and they outnumber the “Match” RGCs by a factor of 10 (Grik4 pool consists of 1030 DREADD AC-driven RGCs, and for Scnn1a the pool has 1151 cells). In summary, we successfully combined near pan-retinal recordings with *post hoc* anatomical characterization and isolated GFP-positive Grik4 and Scnn1a RGCs with increased spiking or bursting activity in the presence of CNO.

In the next step, we identified Grik4 and Scnn1a RGCs according to changes in their spiking pattern following DREADD activation with CNO, and we grouped them into functional clusters based on response similarity.

After the successful registration of spikes with Grik4/Scnn1a GFP RGCs, we grouped these RGCs according to their response preference for stationary (static, full field) and non-stationary (moving, orientations) images. Briefly, cells that were exceeding a threshold value (see Methods) for their direction or orientation selectivity index (DSI and OSI, respectively) were classified as non-stationary.

In the last and final step of our classification protocol, we grouped the registered DREADD RGCs into functional groups according to the nature of their responses to light. We recently described a non-parametric approach for unsupervised RGC classification by using the SPIKE distance (38,39) as a clustering metric (10). In order to use that approach, it is necessary to have a stimulus that elicits responses simultaneously over the entire recording area. Here we used a chirp stimulus inspired from Baden et al (2016) (5) that elicits responses from all RGCs at the same time to pre-sort stationary and non-stationary RGCs (chirp, Fig 6 B, G, top) for SPIKE distance measure and hierarchical agglomerative clustering. We manually validated our detected clusters by grouping several response parameters, e.g. bias index (ON, ON-OFF or OFF) or response duration (transient or sustained), from the chirp and moving bars (for example Fig 6 D-F, I). Manual grouping was only feasible because the expected cluster numbers for Grik4 and Scnn1a were small (respectively 3 and 4 according to our IHC analyses). The pairwise SPIKE distances were determined from all trials of the chirp stimulus and the resulting distance matrix of stationary and non-stationary RGCs was clustered with a hierarchical clustering algorithm followed by the construction of a dendrogram as shown in Jouty et al., 2018 (10). To find the optimal number of clusters for stationary and non-stationary SPIKE distances we used gap statistics (10,40). For stationary and non-stationary Grik4 RGCs, gap statistics estimated three and five response clusters (Fig 6 A), respectively.

**Figure 6:**
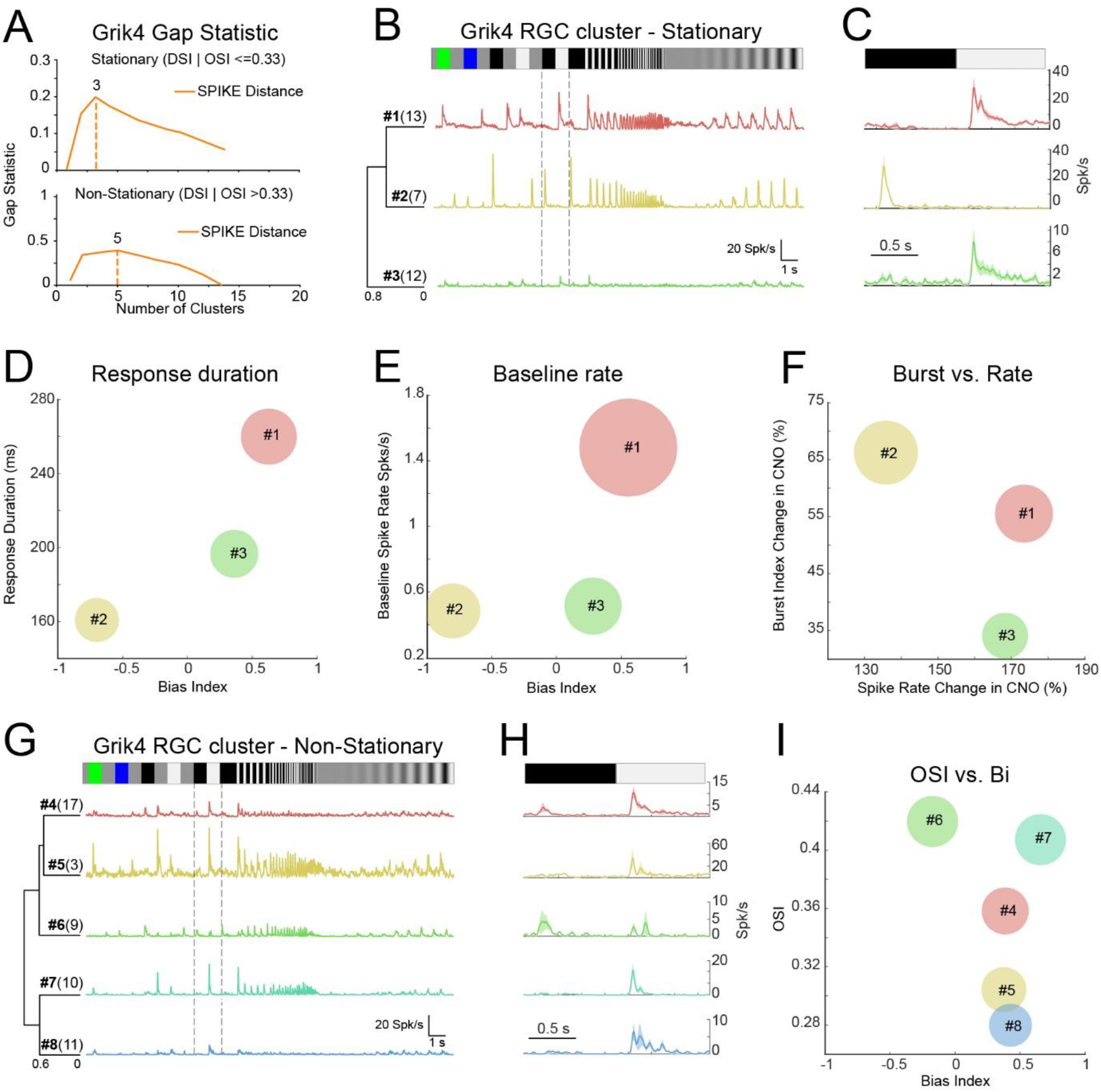
Clustering of Grik4 stationary and non-stationary RGC responses. RGC responses that showed a high spike train similarity for a chirp stimulus (B, F) were grouped together using gap statistics (A). For each RGC of the groups, the PSTH was calculated and the mean Chirp PSTH was plotted (B, G, coloured lines) together with their mean PSTH to a black-white contrast step (C, H). The means for Bias Index (D, E, I), Response Duration (D), Baseline Spiking (E), Burst change (F), Spike Rate change (F) and orientation selectivity index (OSI, I) index were scatter plotted and their standard deviations were used as the circle diameters.

For stationary Grik4, we found ON sustained (#1, n = 13), OFF transient (#2, n = 7) and weak ON sustained (#3, n = 12) response types. To validate these three detected stationary clusters, we plotted the mean bias index against the mean response duration from all RGCs in a cluster (Fig 6 D), revealing three distinct groups that corresponded to the confirmed stationary RGC types. Moreover, spontaneous activity follows very specific patterns for certain RGC types and can therefore help classification. Plotting the mean Bias Index against mean spontaneous firing rate (baseline firing) revealed that the ON sustained cluster #1 (Fig 6 B, E) exhibited a very strong baseline firing rate (Fig 6 E, red bubble). The same plot also shows that the two other RGC types have a moderate baseline firing rate and further confirmed the three well separated RGC types. Whether the spontaneous firing rate becomes more bursting or simply increases monotonously in the presence of CNO can be used to further group RGCs into clusters (Fig 6 F). Interestingly, the effect of CNO for bursting on cluster #3 is minimal (Fig 6 F, green) whereas it is higher for clusters #1 & #2 (Fig 6 F, red & yellow).

Most of the non-stationary RGCs barely responded to the chirp stimulus (Fig 6 G, H) but that response was, albeit modest, reliable and unique for the different clusters. We used the orientation (OS) and direction selectivity (DS) index in addition to the chirp PSTH parameters to define the five non-stationary Grik4 clusters. We found ON-OFF direction selective (DS) (#4, n = 17), ON trans. DS (#5, n = 3), ON-OFF orientation selective (OS) (#6, n = 9), ON trans. OS (#7, n = 10) and ON sust. DS (#8, n = 11) response groups in the pool of non-stationary Grik4 RGCs. All groups were DS; hence we used the OS index vs Bias Index (BI) for our manual validation. All clusters are clearly distinguishable from each other (Fig 6 I). Clusters #4, 5, and 8 have a similar mean BI, but Cluster #7 tends to be marginally more ON and OS (Fig 6 I). Cluster #6 is an OS and ON-OFF cell. In summary (Tab 1), we found three stationary RGCs types and five non-stationary RGC types that share the Grik4 gene pool.

**Table 1.** Summary of Grik4 RGC types with cross-reference to known types

| Grik4 cluster | Baden et al., 2016 | Tran et al., 2019 | Farrow et al., 2013 | Other |
| --- | --- | --- | --- | --- |
| #1 ON sust. | G22, ON sust. |  | PV1 | PixOn, Johnson et al., 2018 |
| #2 OFF trans. | G8, OFF alpha trans | 45 | PV5 | OFF-t, Krieger et al., 2017 |
| #3 ON sust. |  |  |  | Weak ON sust. responses |
| #4 ON-OFF DS | G12, ON-OFF DS | 10, 16, 24 | PV0 | OO-DSGC, Rivlin-Etzion et al., 2011 |
| #5 ON trans. DS | G18, ON trans. (DS) |  | PV2/3 |  |
| #6 ON-OFF OS | G14, (ON)-OFF local OS |  |  |  |
| #7 ON trans. OS | G17, ON local trans OS |  |  |  |
| #8 ON sust. DS | G26, ON sust. DS |  | PV1 |  |

We proceeded in a similar way with stationary and non-stationary registered Scnn1a RGCs (Fig 7). Gap statistics suggested six clusters for stationary and seven for non-stationary clusters (Fig 7 A). The dendrogram of the chirp and black-white contrast PSTH consists of OFF transient (#1, n = 9), ON-OFF (#2, n = 3; #3, n = 25; #4, n = 15), ON sustained (#5, n = 11) and ON transient (#6, n = 18) like responses for stationary Scnn1a RGC types (Fig 7 B, C). Further analysis revealed that the response parameters (Fig 8 D - Bias Index vs Response Duration; Fig 8 F – Burst vs Spike Rate Change) were clearly distinct from each other. At the same time, the distinction is less clear when comparing Baseline vs Bias Index (Fig 7 E). The effect of CNO on the different Scnn1a RGCs was also diverse (Fig 7 F) with cluster #5 barely affected and #2 profoundly affected.

**Figure 7:**
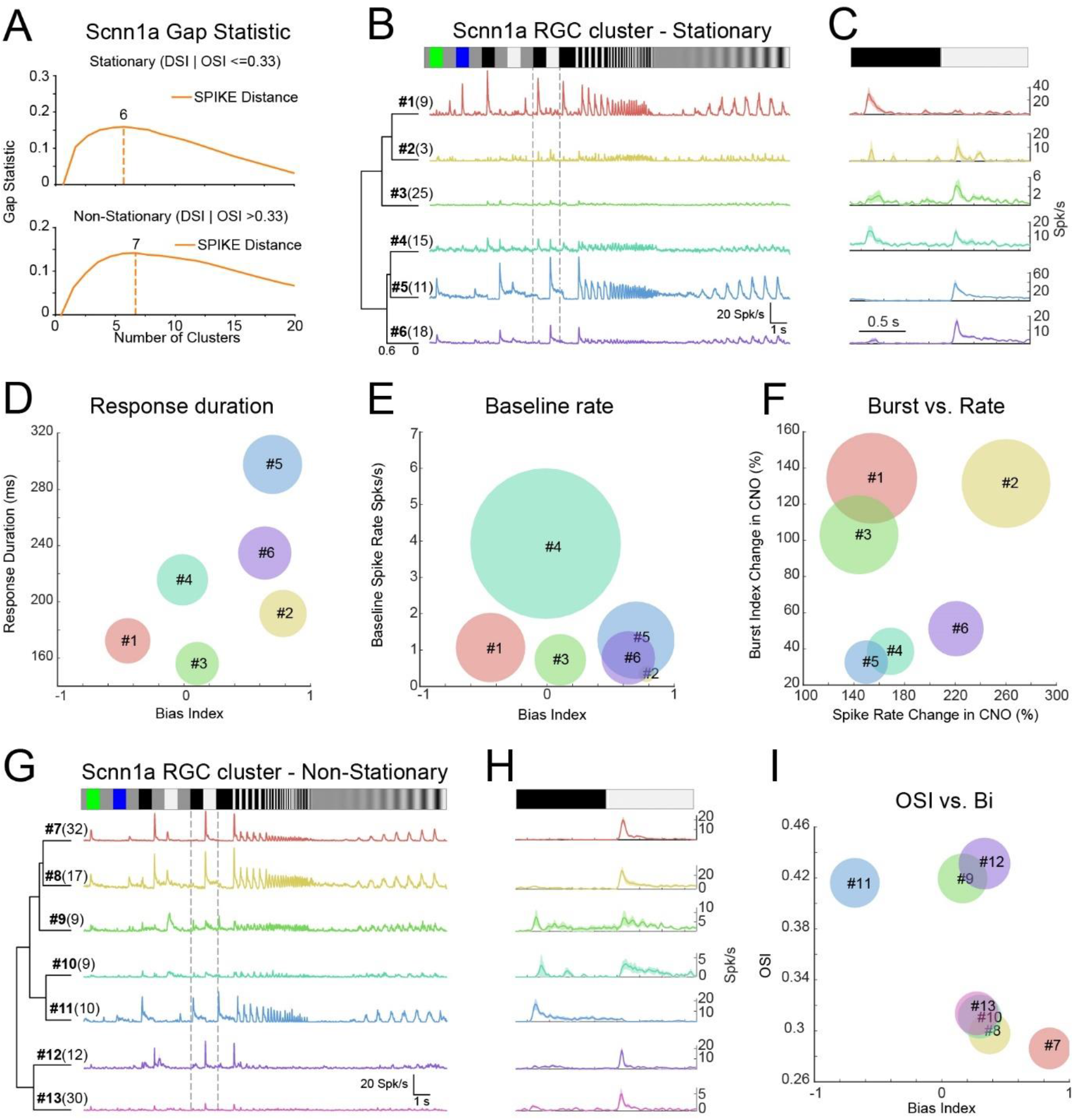
Clustering of Scnn1a stationary and non-stationary RGC responses. RGC responses that showed a high spike train similarity for a chirp stimulus (B, G) and black-white contrast (C, H) were grouped together using gap statistics (A). For each RGC of the groups, the PSTH was calculated and the mean PSTH was plotted (B, C, G, H, colored lines). The means for Bias Index (D, E, F), Response Duration (D), Baseline Spiking (E), Burst change (F), Spike Rate change (F) and orientation selectivity index (OSI, I) index were scatter plotted and their standard deviations were used as the circle diameters.

**Figure 8:**
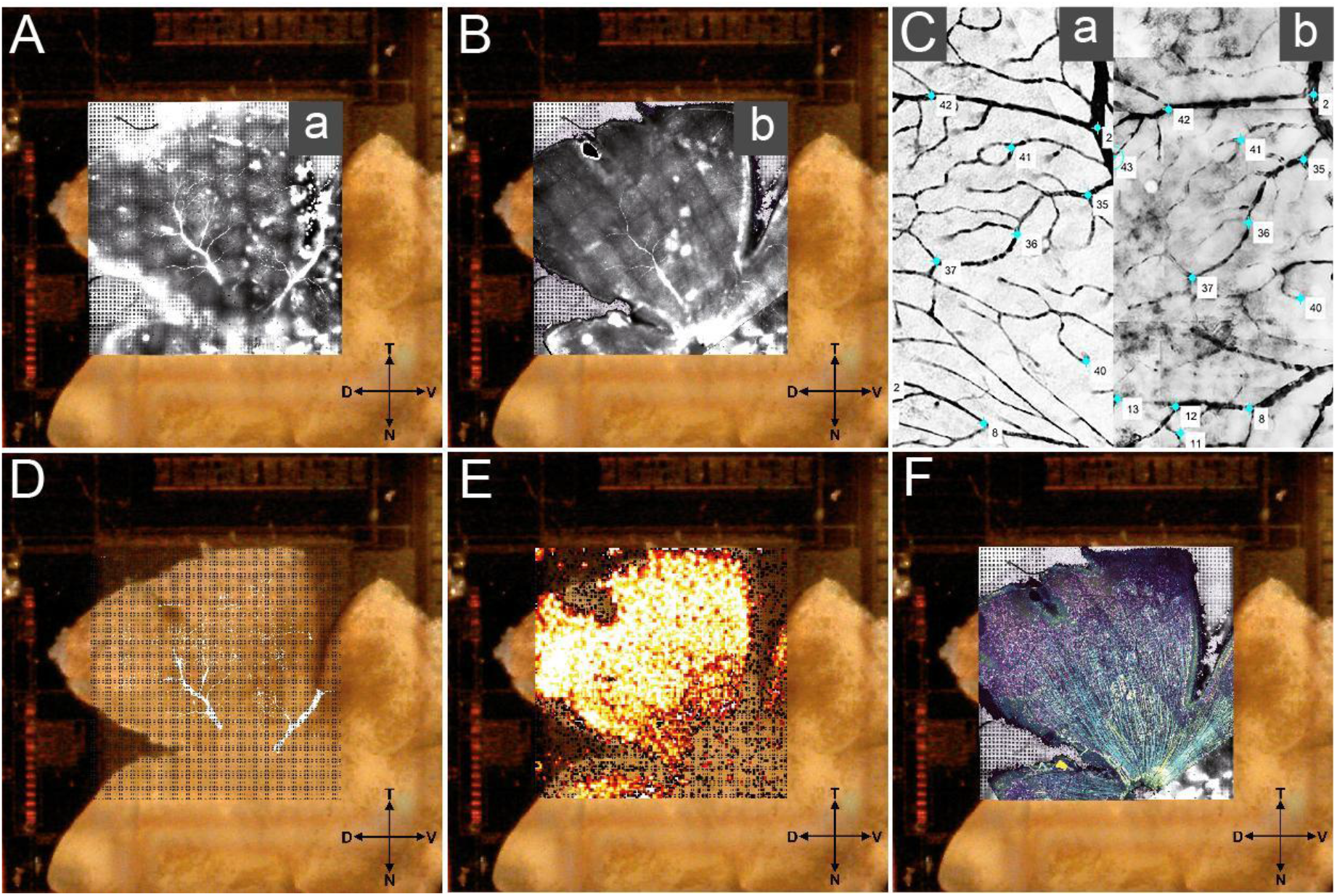
Registering spike location with cell labelling using blood vessel landmarks. Control points in the fixed pre-labelled (A) and moving post-labelled (B) blood vessel image were aligned (C, insets a & b from A and B, respectively) and the resulting transformation matrix (D) was fixed aligned with the spike locations (E) and additional immunostainings (F). D = dorsal; V = ventral; N = nasal; T = temporal; Grid electrode size 2.65 x 2.65 mm

The non-stationary Scnn1a RGCs (Fig 8 F) clustered into ON transient DS (#7, n = 32), ON sustained DS (#8, n = 17), ON-OFF OS (#9, n = 9), ON-OFF DS (#10, n = 9), OFF OS (#11, n = 10), ON OS (#12, n = 12) and ON DS (#13, n = 30) like responses. Although there was some overlap between certain clusters (#8, #10 and #13) when plotting the Bias Index and OS index means (Fig 7 I), their chirp PSTH plots were substantially different, suggesting that these cells do belong to distinct functional groups. Therefore, it is likely that the Scnn1a RGC pool consists of several DS and OS cells which will necessitate further investigation to establish their basic functional differences. In summary (Tab 2), we found a minimum of six stationary and a maximum of seven non-stationary Scnn1a RGC types.

**Table 2.** Summary of Scnn1a RGC types with cross-reference to known types

| Scnn1a cluster | Baden et al., 2016 | Tran et al., 2019 | Farrow et al., 2013 | Other |
| --- | --- | --- | --- | --- |
| #1 OFF trans. | G8, OFF alpha trans | 45 | PV5 | OFF-t, Krieger et al., 2017 |
| #2 ON-OFF |  |  |  | Potentially belongs to #4 |
| #3 ON-OFF |  |  |  | Weak ON-OFF responses |
| #4 ON-OFF | G10, local edge W3 | 6 |  | W3, Zhang et al., 2012 |
| #5 ON sust. | G22, ON sust. |  | PV1 | PixOn, Johnson et al., 2018 |
| #6 ON trans | G18, ON trans. |  | PV2/3 |  |
| #7 ON trans. DS | G16, ON DS trans. |  |  |  |
| #8 ON sust. DS | G25, ON DS sust. 1 |  |  |  |
| #9 ON-OFF OS | G14, (ON)-OFF local OS |  |  |  |
| #10 ON-OFF DS | G12, ON-OFF DS | 10, 16, 24 | PV0 | OO-DSGC, Rivlin-Etzion et al., 2011 |
| #11 OFF OS | G1, OFF local OS |  |  |  |
| #12 ON OS | G17, ON local trans OS |  |  |  |
| #13 ON DS |  |  |  | Weak ON DS responses |

## Discussion

We established a novel approach for RGC classification, based on grouping of cells sharing gene expression and similarities in their light response patterns. A previous approach used pharmacogenetics in combination with MEA recordings for Parvalbumin RGC classification (29). PV expressing cells in the retina are manifold with at least 8 RGC types (24,41) and a distinct number of ACs. Other approaches combined MEA recordings with anatomical imaging (42,43) to unequivocally isolate DREADD ganglion cells. We successfully used excitatory DREADD activation in retinal cells in combination with IHC and large-scale retinal electrophysiology to provide a pan-retinal phenotypical description of Grik4- and Scnn1a-expressing RGCs in the mouse retina. We extended the pre-existing list of Grik4 expressing RGC types and, for the first time, we provide a functional description of Scnn1a-expressing RGCs. We also showed that certain types of ACs express Grik4 and Scnn1a and that exciting these cells with DREADD activation leads to changes in firing frequency in many undefined RGCs. We used DREADD instead of the more common optogenetic technique because the retina is light sensitive and we don’t want to use strong lights that interfere with our stimuli. Our approach is not restricted to retinal cells but is widely applicable to other neurons from other brain regions e.g. cortical slices. It is a scalable multimodal approach and can provide fast grouping of large cohorts of neurons with similar gene expression. Additionally, we provide a complete framework to combine high-density MEA recordings with *post hoc* cell labelling.

Grik4 expression in the retina has been described (13,43–46) but there is functional characterization only for two RGC types (45,46). Johnson et al., 2018 (46) reported that the Grik4-expressing PixON RGC type is an ON sustained type and has strong spontaneous activity. Our Grik4 cluster #1 from the stationary RGCs (Tab 1) has the same key phenotypic features - sustained responses and strong spontaneous activity. Cluster #1 also matches with the group (G) 22 from Baden et al., 2016 (5) and PV1 from Farrow et al., 2013 (41). Rivlin-Etzion et al., 2011 (45) reported a Grik4 positive ON-OFF DS RGC type. Such a response type is found in our non-stationary Grik4 cluster #4 which corresponds to G12 (Baden et al., 2016), dorsal-temporal clusters 10, 16, 24 in Tran et al., 2019 and PV0 in Farrow et al., 2013. Our novel classification approach was able to match two previously reported Grik4 response types, and such similarities provide solid validation for the methodology itself. Further, Grik4 stationary cluster #2 potentially resembles G8 (OFF alpha transient) which is further described in Krieger et al., 2017 (47) and in Tran et al., 2019 (cluster 45). The other Grik4 clusters with their cross-reference match are listed in tab 1.

For Scnn1a, we found six stationary and seven non-stationary RGC response groups (Tab 2). Interestingly, the Scnn1a cluster #5 shows remarkable similarity with the PixON characteristics described earlier - sustained responses and high baseline firing. Stationary Scnn1a cluster #1 seems to be an OFF alpha transient type (Krieger et al., 2017). Last, cluster #10 is potentially the ON-OFF DS RGC type reported by Rivlin-Etzion et al., 2011. There are no reports about Grik4 and Scnn1a co-expressing cells but it is not uncommon that certain RGC types share the same genes e.g. *Foxp* AND *Brn3* (30) or *Pvalb* AND *Calb2* (27,30,31). In summary, our approach is able to group RGC response types recorded across large MEA recordings into established response clusters and finds correlates in previous reports (tab 1, 2). However, our approach yields a large number of clusters for each gene pool, demonstrating that it is a powerful classification tool.

Grik4 clusters #1, #2 #4, #5, #8 and Scnn1a clusters #1, #5, #6, #10 are parvalbumin positive (5,14,41,47) and in line with our IHC results that revealed many Grik4 and Scnn1a and parvalbumin expressing RGCs. Baden et al. (2016) found many more parvalbumin-expressing RGC types than the traditionally known 8 types from anatomical studies (24). We suspect that there is some overlap between Grik4 and yet undescribed parvalbumin, potentially also calretinin RGC types. We currently investigate the functional features of parvalbumin and calretinin co-expressing RGCs using a similar methodology (31). Lee et al., 2010 (25) reported calretinin and parvalbumin co-expression in a subset of RGC types that would correlate to our Grik4 clusters #4, #5 and #8, as well as Scnn1a clusters #5, #6, #8 and #10. Here our classification results come full circle and match with the parvalbumin and calretinin IHC results where we find all combinations.

DREADDs were also expressed in one or several yet undescribed AC types. For Scnn1a, we found few GABAergic ACs (not shown) but most GFP-positive ACs were not characterized further. For Grik4, we did not find any GABAergic ACs, nor were we able to define the AC type with our experimental means. The net effect of CNO induced activity in DREADD ACs can have many forms leading to a priori undetermined situations. For example, it is known that cascade of inhibition can result in excitation (e.g. push-pull effect) (48). A possible solution to this problem relies on a quantitative analysis requiring considering the factors constraining individual cell responses without and with CNO, and the network connectivity. It is possible to propose a map of CNO induced scenarios in simple situations, with a suitable space of relevant biophysical parameters. These questions will be addressed on modelling and mathematical grounds in a forthcoming paper.

Finding DREADD expression in ACs was initially challenging. We managed to circumvent the interference of DREADD activated ACs by combining MEA recordings with cell labelling, which renders experiments much more complex. Our experimental design was based on Cre-Lox recombination, resulting in DREADD expression in all cells with the same promoter gene in the organism. In future work, DREADD expression should be targeted only to RGCs via viral transfection in order to avoid such strong side effects (49). Blocking AC input onto postsynaptic targets (bipolar cells or RGCs) would be another solution but has a big caveat: such disinhibition leads to massive increase in activity levels in all RGCs (not shown), hence masking RGC specific DREADD activation in specific subgroups where DREADDs are directly expressed on RGCs. Alternatively, using intracellular recordings from single cells could exclusively target Grik4 and Scnn1a RGCs based on GFP expression, and DREADD activation with CNO would be redundant. Such an approach would provide unequivocal recordings from RGCs, but at the same time, the throughput would be extremely low, and all network information (provided by recording simultaneously from many homotypic RGCs) would be lost (looking at this information will be the subject of a separate publication). Lastly, it is known that the Cre-lox recombination has disadvantages (50) and sometimes leads to off-target expression. We followed the breeding advice from the manufacturer and used animals from different breeding pairs.

In summary, our approach is able to quickly isolate large numbers of neurons at once while retaining all network information. This approach does not have to be restricted to DREADD technology, and our framework can be used also only with cell labelling (see also (31)).

## Methods

### Animals and retina preparation

All experimental procedures were approved by the ethics committee at Newcastle University and carried out in accordance with the guidelines of the UK Home Office, under control of the Animals (Scientific Procedures) Act 1986. Grik4 (C57BL/6-Tg(Grik4-cre)G32-4Stl/J, the Jackson Laboratory, MA, JAX Stock No: 006474) and Scnn1a (B6;C3-Tg(Scnn1a-cre)3Aibs/J, JAX Stock No: 009613) mice were cross-bred with Gq-DREADD mice (B6N;129-Tg(CAG-CHRM3*,-mCitrine)1Ute/J, JAX Stock No: 026220) to generate a strain of mice with the excitatory Gq-DREADD expressed in Grik4 and Scnn1a expressing cells (Grik4-DREADD and Scnn1a-DREADD, respectively). In addition, we crossbred the Grik4 and Scnn1a lines with an inhibitory DREADD (B6.129-Gt(ROSA)26Sortm1(CAG-CHRM4*,-mCitrine)Ute/J, JAX Stock No: 026219) but the effect on RGC firing rate was negligible and the litters were used only for immunofluorescence studies. Male and female wild-type, Grik4-DREADD and Scnn1a-DREADD mice, housed under a 12-hour light-dark cycle and aged between postnatal days (P) 53-148 were used for the experiments. Mice were dark-adapted overnight and killed by cervical dislocation. Eyes were enucleated, and following removal of the cornea, lens, and vitreous body, they were placed in artificial cerebrospinal fluid (aCSF) containing the following (in mM): 118 NaCl, 25 NaHCO_3_, 1 NaH_2_PO_4_, 3 KCl, 1 MgCl_2_, 2 CaCl_2_, 10 glucose, and 0.5 l-Glutamine, equilibrated with 95% O_2_ and 5% CO_2_. The ventral and dorsal orientation was marked after enucleation. The retina was isolated from the eye cup and flattened for MEA recordings. For vertical cryosections, mouse eyecups were fixed in 4% paraformaldehyde (PFA; Alfa Aesar, MA) in 0.1 M phosphate buffer solution (PBS) for 2 x 20 minutes at room temperature and washed with PBS several times. For whole mounts, retinas were isolated from the eye cup and mounted on nitrocellulose paper (Sartorius, Germany) and transferred to 4% PFA in PBS (2 x 20 min), rinsed in PBS and prepared for further procedures. All procedures involving live animals and retinas were performed in dim red light and the room was maintained in darkness throughout the experiment.

### Immunohistochemistry and image acquisition

After enucleation and fixation, the retinal tissue was processed in different ways for vertical and whole mount IHC. For vertical sections, eyecups were cryoprotected in 30% sucrose in PBS overnight at 4°C and embedded in OCT Tissue TeK (Sakura, NL) at −20°C on the following day. Vertical sections (15-20 μm) were cut on a OTF5000 cryostat (Bright Instruments, UK) and collected on Superfrost microscope slides (Thermo Fisher). Vertical sections and whole mounts were blocked with 10% normal goat serum (NGS) and/or 10% normal donkey serum (NDS) in PBS for at least 30 minutes at room temperature.

After the blocking procedure and a short rinse in PBS, vertical sections were incubated with primary antibodies in 5% NGS (and/or NDS) + 1% Triton X-100 + PBS overnight at 4°C. Whole mounts were incubated free-floating with primary antibodies in 5% NGS (and/or NDS) + 1% Triton X-100 + PBS for 4-5 days at 4°C. Incubation with secondary antibodies in 1% Triton X-100 in PBS was carried out for 2 hours at room temperature for vertical sections or overnight at 4°C for whole mounts. Details of the primary antibodies are as follows: anti-GFP (chicken, Abcam 13970, 1:500-1000), anti-RBPMS (rabbit, Phosphosolutions 1830, 1:1000), anti-GABA (mouse, Sigma Aldrich A0310, 1:1000), anti-Calretinin (mouse, Swant 6B3, 1:1000) and anti-Parvalbumin (rabbit, Swant PV27, 1:1000). Secondary antibodies are as follows (all concentrations 1:500): goat anti-chicken CF488, goat anti-rabbit Alexa568/Alexa647 and donkey anti-mouse Alexa647. After washing several times with PBS, sections and whole mounts were mounted in Vectashield (Vector Laboratories, UK). All incubations and washing procedures were performed in the dark.

Images were captured using either a Zeiss Axio Imager upright microscope with Apotome structured illumination fluorescence (using 20x/40x air objectives) or a Zeiss LSM800 confocal microscope with 40x oil objective (Zeiss, Germany) operated with Zen software. Whole mount images were stitched together using Zen software. All other image post processing was done with Fiji (https://fiji.sc), Adobe Photoshop (Adobe, CA) and MATLAB (Mathworks, MA). The steps between single sections of confocal stacks were not exceeding 1 μm and 3 - 5 sections were superimposed with Fiji for presentation. For the cell density maps, cells were manually counted using the “Cell Counter” plugin in Fiji and a bivariate histogram (MATLAB *hist3*, bin size 60 x 60 μm) was calculated for the cell densities. For visualizing purposes, a 2-D Gaussian filtering was applied (MATLAB *imgaussfilt*, sigma 3).

### Large-scale, high-density multielectrode array recordings and light stimulation

Recordings were performed on the BioCamX platform with high-density-multielectrode array (HD-MEA) Arena chips (3Brain GmbH, Lanquart, Switzerland), integrating 4096 square microelectrodes in a 2.67 x 2.67 mm area and aligned in a square grid with 42 μM spacing. The isolated retina was placed, RGC layer facing down, onto the MEA chip and flattened by placing a small piece of translucent polyester membrane filter (Sterlitech Corp., Kent, WA, USA) on the retina followed by a home-made anchor. Retinas were maintained at 33°C using an in-line heater (Warner Instruments LLC, Hamden, CT, USA) and continuously perfused using a peristaltic pump (~1 ml min-1). Retinas were left to settle on the MEA for at least 2 hours before recording. The platform records at a sampling rate of ~18 kHz/electrode when using the full 64×64 array. Recordings are filtered at 50Hz high-pass filter using BrainWaveX software (3Brain) and stored in hdf5-based data format. Spikes were detected and sorted using Herdingspikes2 (https://github.com/mhhennig/HS2) as in (16). Briefly, spikes were first detected as threshold crossings individually on each channel, and then merged into unique events based on spatial and temporal proximity. For each detected spike, a location was estimated based on the signal centre of mass. Spike sorting was performed by clustering all events using a feature vector consisting of the locations and the first two principal components of the largest waveform.

Light stimuli were projected onto the retina as described elsewhere (17). Briefly, the projector irradiance was attenuated using neutral density filters to mesopic light levels (white 4 μW/cm^2^). For stimuli we used a full field ‘chirp’ stimulus consisting of various 1-sec contrast steps, increasing frequency (1-15Hz) and contrast modulations (1-93 Michelson contrast) which was repeated 5 times. We also used random black and white moving bars (width 100 μM, 12 directions (30° separation)), 800 μm/s, and the whole sequence was repeated 5 times. For the chirp and motion stimuli, we estimated each unit’s instantaneous firing rate for the different stimuli by convolving its spike train with a Gaussian kernel smoothing function (standard deviation (SD) = 25 ms). We then averaged the trials and extracted several features including the Bias Index and the response duration (see (16)). On average, recording from one retina yields light responses from hundreds to thousands of individual RGCs.

### Registering RGC activity with IHC - blood vessels as reference marker

To register RGC activity with IHC, we used pre- and post-labelled blood vessels as reference markers (Fig. 1). For pre-labelled blood vessel staining, the eye cup was incubated for 1 hr in aCSF + 5 μM Sulforhodamine 101 (SR101, Sigma Aldrich, MO) and afterwards transferred to aCSF + 0.06 μM SR101 for MEA recording procedures. After recording, the weight and the polyester membrane were carefully removed to expose the retina, and the MEA well rinsed with aCSF. If the blood vessels were not visible, a 50% Optiprep (Sigma Aldrich, MO) /50% aCSF solution was used for rinsing and further clearing. The MEA chip was mounted on a microscope stage (Olympus AX70, CoolLED pE fluorescence) while continuing rinsing every 2-3 min with oxygenated aCSF to ensure the tissue remains healthy throughout the procedure. The best stained blood vessel layer (either superficial or deep vascular plexus) was meander-like imaged (Fig. 8 A). For every captured blood vessel, images were acquired in the same location all the way down to the electrodes as well. Neighbouring images were manually stitched together in Adobe Photoshop (Adobe, CA) later. The retina was then carefully removed from the MEA, flattened on a nitrocellulose membrane for IHC procedure. Blood vessel staining was enhanced by using an anti-mouse secondary antibody in the same excitation spectrum (normally Alexa568) and blood vessels were imaged again (same plexus as while the retina was imaged on the MEA (Fig. 8 B). Both blood vessel staining images, pre- and post-tissue fixation were correlated with each other using the Control Point Selection Tool in Matlab (Fig. 8 C). Briefly, the images taken in the live retina on the MEA were used as the fixed (reference) image and images acquired post fixation as the moving image. Minimum 50 reference points were picked for the fixed and moving images to create a geometric transformation (Matlab *fitgeotrans, lwm, 50*). That geometric transformation was applied to the GFP image. A stack (hereafter called MEA images) with the MEA electrode image, blood vessels pre- and post-fixation and GFP image was created in Adobe Photoshop (Fig. 8 F).

### Registering RGC activity with IHC - Grik4/Scnn1a DREADD RGC identification

A full protocol for the next steps with code and example data will be made publically available on our GitHub repository (https://github.com/GerritHilgen/DREADD_RGCclass). Briefly, units that did show a significant change in spontaneous firing rate (sampled during at least 5 min) before and after adding CNO to the chamber were selected for Grik4/Scnn1a RGC identification and response clustering. Units with >50% change in firing rate in CNO were considered as potential Grik4/Scnn1a candidates. We also used the Burst Index (33) to look at potential changes in activity levels induced by CNO, with potential Grik4/Scnn1a cells having changes in their Burst Index exceeding > 50%. This is followed by correlating the physical position of these identified RGCs with micrographs of DREADD-GFP expressing cells in the GCL. During the spike sorting process with Herdingspikes2, the physical x and y spike coordinates for each detected spike from a single RGC unit is calculated. The spike location coordinates are given for a 64 x64 grid (the electrode layout of our MEA). GFP images were transformed to 8-bit images, a threshold (Matlab *max entropy)* was applied, potential gaps filled (Matlab *imfill ‘holes’*) and binarized (Matlab *bwareaopen 50*). Binary images were analyzed (Matlab *regionprops*), centroids and diameters for every potential GFP blob were calculated and the pairwise distance between centroids and spike locations was measured for all combinations. Both, spike centres and GFP cells, were overlayed.

### Registering RGC activity with IHC - Grik4/Scnn1a response clustering

All potential Grik4/Scnn1a RGC units linked to a GFP blob were further analyzed and grouped according to their response to a chirp stimulus. NB it is a two-factor isolation: 1: the RGC has to have a change of activity of > 50% and has to be within 60 μm of a DREADD GFP cell. But before that grouping, cells were pre-classified into motion sensitive (responding preferably to either direction or orientation of motion) or cells that respond best to static stimuli (full field, chirp). We calculated the direction (DSI) and orientation (OSI) selectivity index as described elsewhere and used a cut-off value of 0.33 (16). The trial variability and signal to noise ratio was calculated (16) for each stationary/non-stationary RGC, and units with a value below the 25^th^ percentile were discarded. The chirp responses of the remaining Grik4 and Scnn1a stationary/non-stationary RGC units were used to establish response groups. We used a similar approach to calculate the SPIKE distance as described in our earlier work (10). Briefly, the SPIKE distances were computed using the open-source package PYSPIKE (39). The pairwise distances between two units were determined by computing the pairwise distances of all trials of the chirp stimulus. The resulting distance matrix was then clustered with a hierarchical clustering algorithm (Python, *Scipy library dendrogram*). Matlab and Python scripts and other extended material will be made available via GitHub.

## Acknowledgments

This project was funded by the Leverhulme Trust (RPG-2016-315 to ES and BC), by Newcastle University (Faculty of Medical Sciences Graduate School and Pro-Vice Chancellor Discretionary Fund). We thank Matthias Hennig for help with the SPIKE distance calculations and Chris Williams who worked on related, unpublished aspects of the project. We also thank the Bioimaging Unit at Newcastle University for providing excellent service and help for this project.

## Authors’ contributions

Conceptualization: G.H., E.S., B.C.; Software: G.H. Methodology: G.H., E.S.; Formal Analysis: GH., Investigation, G.H., V.K., E.K.; Writing - Original Draft: G.H., E.S., B.C., V.K., E.K.; Project administration: E.S., B.C.; Funding Acquisition: E.S., B.C.

## Notes

Financial interests or conflicts of interest: The authors declare no competing financial interests

### Competing Interest Statement

The authors have declared no competing interest.

### Summary of Updates

We got substantial peer review feedback and integrated the changes in the new manuscript - new figures 2,4 and 5 - example data set and code tpo play with (in text link) - revised table 1 and 2

